# An engineered stable mini-protein to plug SARS-Cov-2 Spikes

**DOI:** 10.1101/2020.04.29.067728

**Authors:** Maria Romano, Alessia Ruggiero, Flavia Squeglia, Rita Berisio

## Abstract

The novel betacoronavirus SARS-CoV-2 is the etiological agent of the current pandemic COVID-19. Like other coronaviruses, this novel virus relies on the surface Spike glycoprotein to access the host cells, mainly through the interaction of its Receptor Binding Domain (RBD) with the human angiotensin-converting enzyme 2 (ACE2). Therefore, molecular entities able to interfere with binding of the SARS-CoV-2 Spike protein to ACE2 have a great potential to inhibit viral entry. Starting from the available structural data on the interaction between SARS-CoV-2 Spike protein and the host ACE2 receptor, we here engineered a mini-protein with the aim of creating a soluble and stable Spike interactor. This mini-protein, which was recombinantly produced in high yields, possesses a stable α helical conformation and is able to interact with the RBD of glycosylated Spike protein from SARS-CoV-2 with nanomolar affinity, as measured by microscale thermophoresis. By plugging the Spike protein, our mini-protein constitutes a valid tool for the development of treatments against different types of coronavirus.

## Introduction

The novel coronavirus SARS-CoV-2 has spread widely and rapidly since it was first identified in Wuhan, China in December 2019 [1-3]. Its associated disease, COVID-19, causes severe respiratory difficulties, with aged patients at higher risk of mortality [1]. At the time of writing, over 3.000.000 confirmed cases and over 200.000 deaths have been registered worldwide. Given the dramatic public health emergency, there is a strong and urgent need for new antiviral agents to block human-to-human transmission and to treat infected patients.

Like other coronaviruses, SARS-CoV-2 makes use of a densely glycosylated Spike (S) protein to gain access into host cells [4,5]. The S protein forms homotrimers protruding from the viral envelope and binds with high affinity to the host receptor ACE2 (angiotensin-converting enzyme 2), mainly expressed by epithelial cells of the respiratory tract [6]. Recently, another human receptor, CD147, has been identified as a possible route of viral entrance, again mediated by the S protein [7]. CD147 is also known as Basigin or EMMPRIN, and is a transmembrane glycoprotein that belongs to the immunoglobulin superfamily involved in many processes, including tumor development, plasmodium invasion and virus infection [7]. After attachment, the human transmembrane protease serine 2 (TMPRSS2) cleaves and activates the S protein, thus allowing SARS-CoV-2 to enter the host cells [8]. Compared to SARS-CoV, an additional protease, possibly furin, is likely involved in priming of the SARS-Cov-2, since the S protein of SARS-CoV-2 contains four redundant furin cut Pro-Arg-Arg-Ala motifs, which are absent in SARS-CoV [6]. The S protein contains two subunits, S1 and S2. Of these, S1 comprises the receptor binding domain (RBD), which is responsible for recognizing and binding the cell surface receptor ACE2 [5]. Being essential for infection, the S protein is a promising target for antibodies and vaccines [9]. Indeed, preventing attachment of the S protein to either the ACE2 or the CD147 receptors would hamper infection at the early viral entry step. Based on the structural information available on the complex between the S protein and the ACE2 receptor [10], we engineered a miniaturised mimic of ACE2, here named Spikeplug. This mini-protein, which we produced in high yields, is highly soluble, conformationally stable and displays nanomolar binding affinity with the S protein of SARS-CoV-2. Given its properties, Spikeplug is a promising molecule for the development of treatments based on inhibition of viral entry for different types of coronavirus.

## Results

### Spike interactor design and synthesis

We used all available structural information on the interactions between S protein and the human ACE2 receptor to design Spike interactors. An analysis of the structure of SARS-CoV-2 complex with ACE2, reveals that most hydrogen bonds between protein S and ACE2 involve the helix H1 (residues 20-52) and the C-terminal part of helix H2 (residues 56-82) of ACE2 [10]. Outside this region, an important interaction between the S protein and ACE2 is mediated by Lys353 of ACE2. Indeed, the presence of histidine in the place of lysine at position 353 of rat ACE2 disfavours viral binding [11].

Starting from the region H1-H2 of ACE2 as a basic scaffold for the design of a Spike interactor, we included a number of mutations to increase its stability and solubility and with the aim to compensate for the missing interactions mediated by residues outside helices H1 and H2. An important issue with peptide or small protein design is their conformational stability, since peptides are usually not folded in solution. Therefore, to stabilise this scaffold, we included an extra helix, H3 (residues 91-101) which naturally caps helices H1 and H2 in the ACE2 structure, through a cluster of hydrophobic interactions involving Val92, Leu97 and Leu100 of H3, Phe28, Leu29, Phe32 of H1 and Ala80 of H2, therefore holding together the two helices H1 and H2 (Figure 1). In addition, we noticed that Glu37 is important to stabilise the structure of the ACE2 receptor by forming a salt bridge interaction with Arg393 and a hydrogen bond with the backbone N of Lys353 (Figure 2A). However, this residue forms a weak hydrogen bond with Tyr505, which is considered outside the Receptor Binding Motif (RBM) of the RBD domain (Figure 2A), in the ACE2 complex with the RBD of the S protein from SARS-CoV-2 [10], while it is not engaged in any contact in the complex of ACE2 with RBD of the S protein of SARS-CoV (74% homologous to SARS-CoV-2 and 83% on the RBD domain) [11]. Therefore, we mutated Glu37 to an arginine residue and energy minimised using GROMACS [12]. As shown in Figure 2A, Arg37 side chain can form hydrogen bonds with the main chain of Gly496 of the S protein (Figure 2A). As such, the conformation of Arg37, also stabilised by a salt bridge with Asp38, mimics the interactions with the S protein mediated by Lys353 of ACE2 (Figure 2A) [10,11]. We further engineered this variant by including two mutations, Leu 91 to Gln and Leu 95 to Gln, to improve protein solubility by replacing hydrophobic with hydrophilic residues, and hamper aggregation in solution. In ACE2, these two residues are located on the opposite side of the S-binding region and form hydrophobic interactions with the core of ACE2 to stabilise it (Figure 2B,C). Their mutation to glutamine residues was also motivated by the characteristics of glutamine to further stabilize α helix structures. This recombinant mini-protein, here named Spikeplug (Table 1, Figure 2C), was successfully over-expressed in *E*.*coli*, resulting in a high yield, of 70 mg of pure protein from one litre of bacterial culture.

**Table 1.**
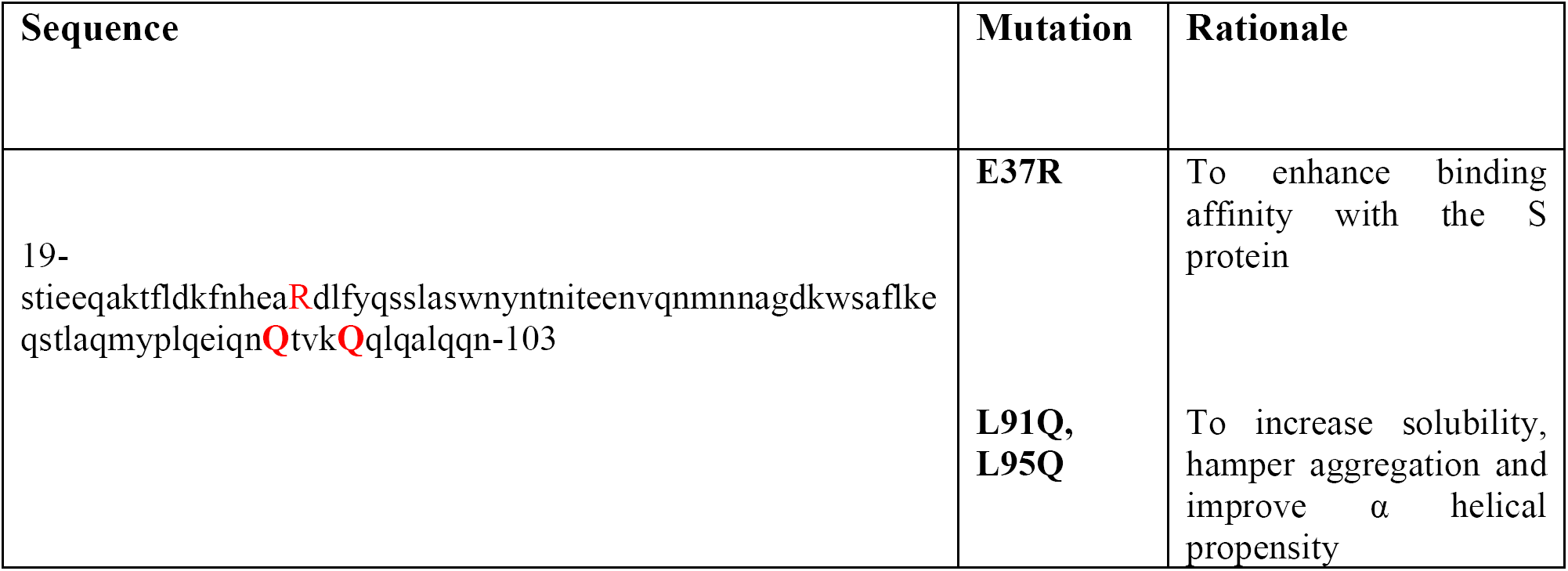
Sequence of Spikeplug. Mutations are drawn in capital red letters.

| Sequence | Mutation | Rationale |
| --- | --- | --- |
| 19-stieeqaktfldkfnhea**R**dlfyqsslaswnyntniteenvqnmnnagdkwsaflke<br>qstlaqmyplqeiqn**Q**tvk**Q**qlqalqqn-103 | **E37R**<br>**L91Q,**<br>**L95Q** | To enhance binding affinity with the S protein<br>To increase solubility, hamper aggregation and improve $\alpha$ helical propensity |

**Figure 1.**
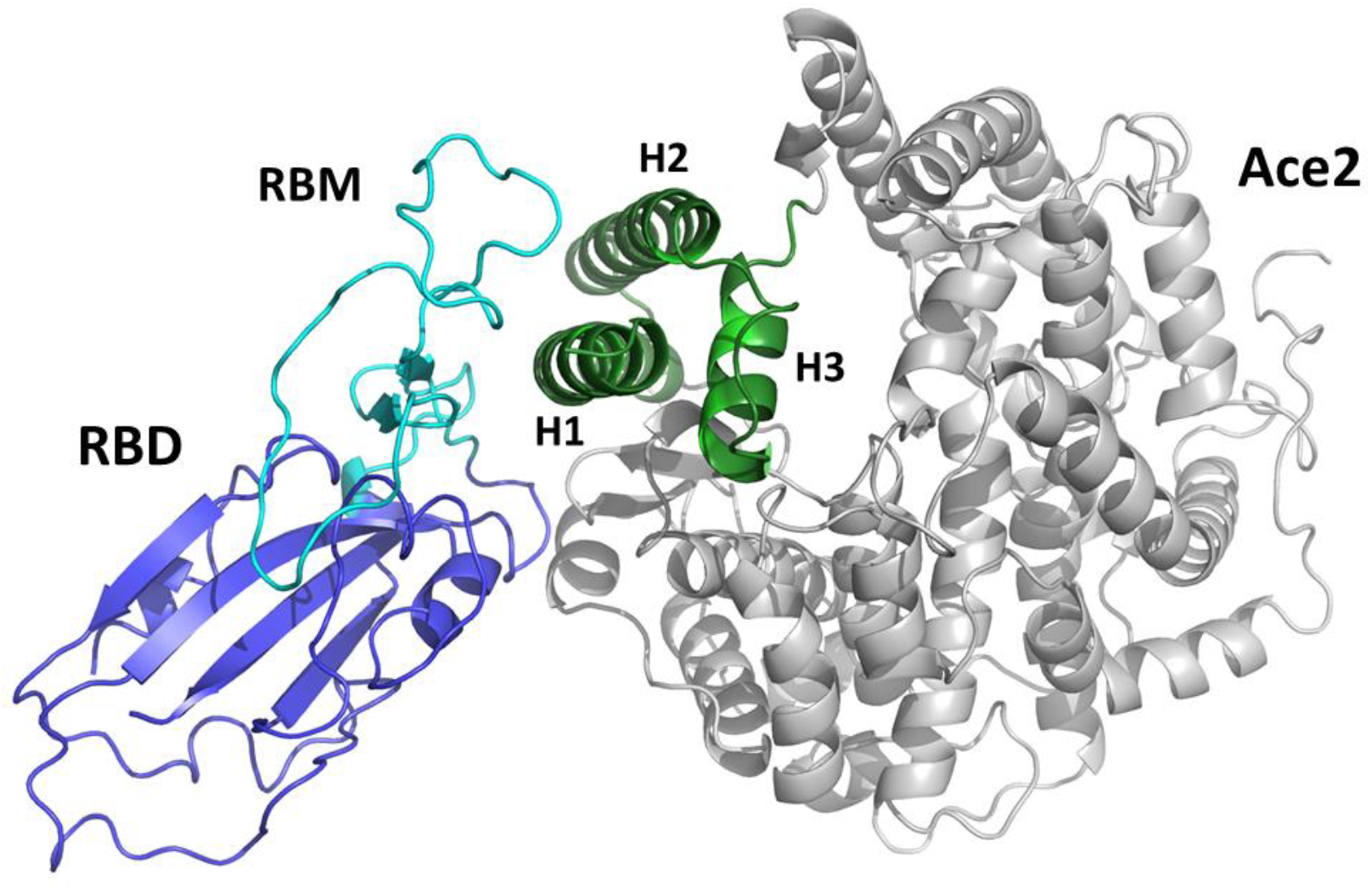
Cartoon representation of the complex between the receptor binding (RBD) domain of SARS-CoV-2 Spike protein (blue/cyan) and the human ACE2 receptor (grey/green). The receptor binding motif (RBM) is drawn in cyan. The green portion of the ACE2 domain including helices H1, H2 and H3 is drawn in green.

**Figure 2.**
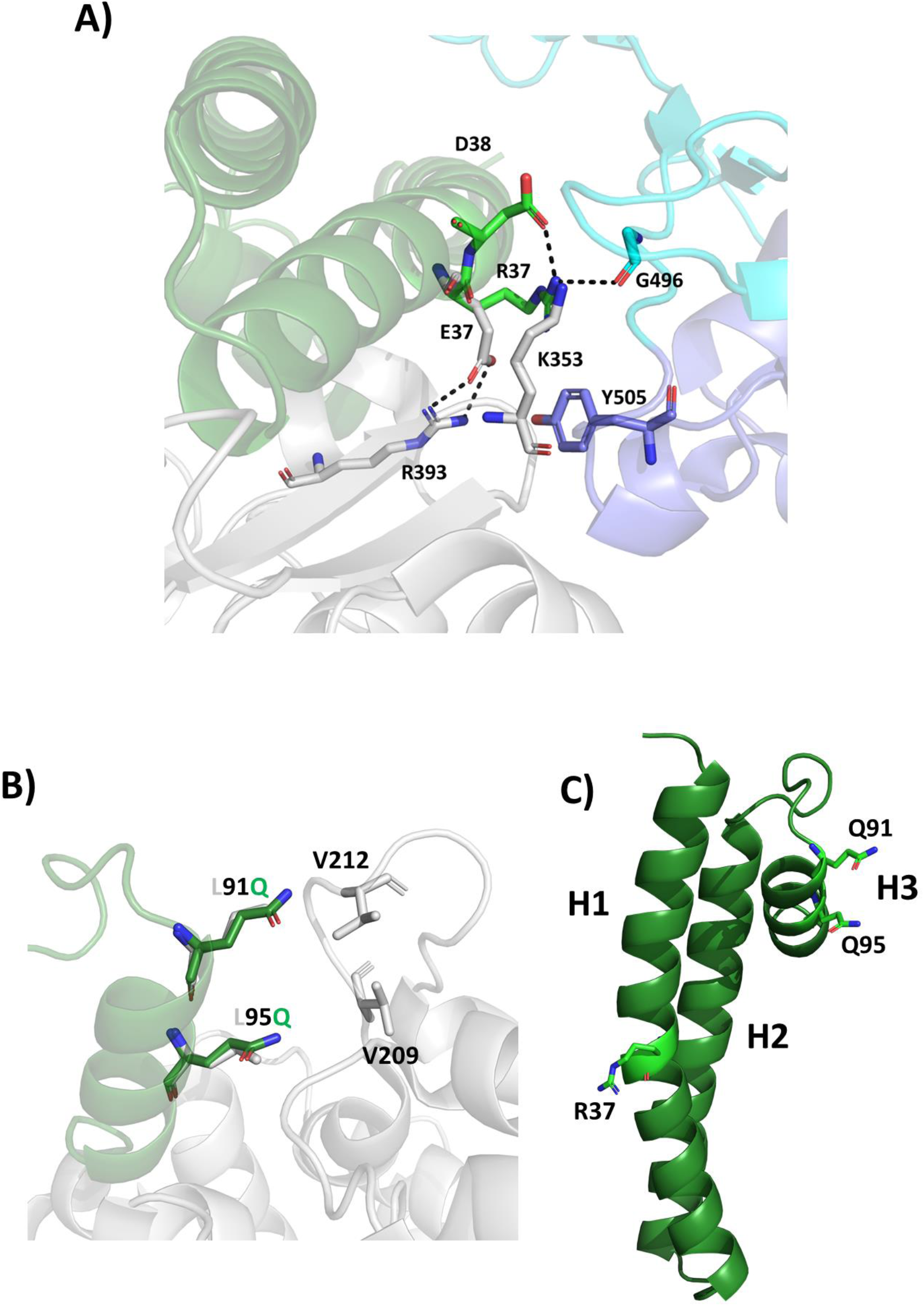
Structural basis of engineered mutations. Cartoon representations showing (A) the superposition of Spikeplug (green) on the the complex between ACE2 (grey) and protein S (cyan-blue). In the structure of ACE2-S, Glu37 of ACE2 forms an intramolecular salt bridge with Arg393 and the backbone of Lys353. The side chain of Lys353 forms hydrogen bonds with Asp38 and the backbone carbonyl oxygen of Gly496 of the S protein. In the minimised Spikeplug-S complex, Arg37 replaces Lys353 in forming hydrogen bonds with both Asp38 and Gly496 of the S protein. (B) the superposition of Spikeplug (green) on ACE2 (grey). Leu 91 and Leu95, involved in hydrohobic interactions with V209 and V212 in the ACE2 structure, were mutated to glutamine residues in Spikeplug. (C) the structure of Spikeplug, mutates residues are shown in stick.

### Structural stability and binding studies

Using far-UV CD spectroscopy, we observed that the spectrum of Spikeplug is typical of a well-structured fold with a high α-helical content (Figure 3A), with typical minima at 208 and 222 nm (Figure 3). Spectra also showed that folding is fully reversible, with the CD spectrum after refolding fully superimposable to that recorded at 4°C (Figure 3A). To investigate the heat-induced changes in the protein secondary structure, thermal unfolding curves were recorded by following the CD signal at 208 nm as a function of temperature, thus providing a melting temperature Tm of 37°C (Figure 3B). Microscale thermophoresis (MST) analysis was performed to measure the binding affinity of Spikeplug to the glycosylated S protein of SARS-CoV-2. MST measurements showed nanomolar binding affinity of Spikeplug to the Spike RBD domain, with a K_D_ of 40 nM (Figure 4).

**Figure 3.**
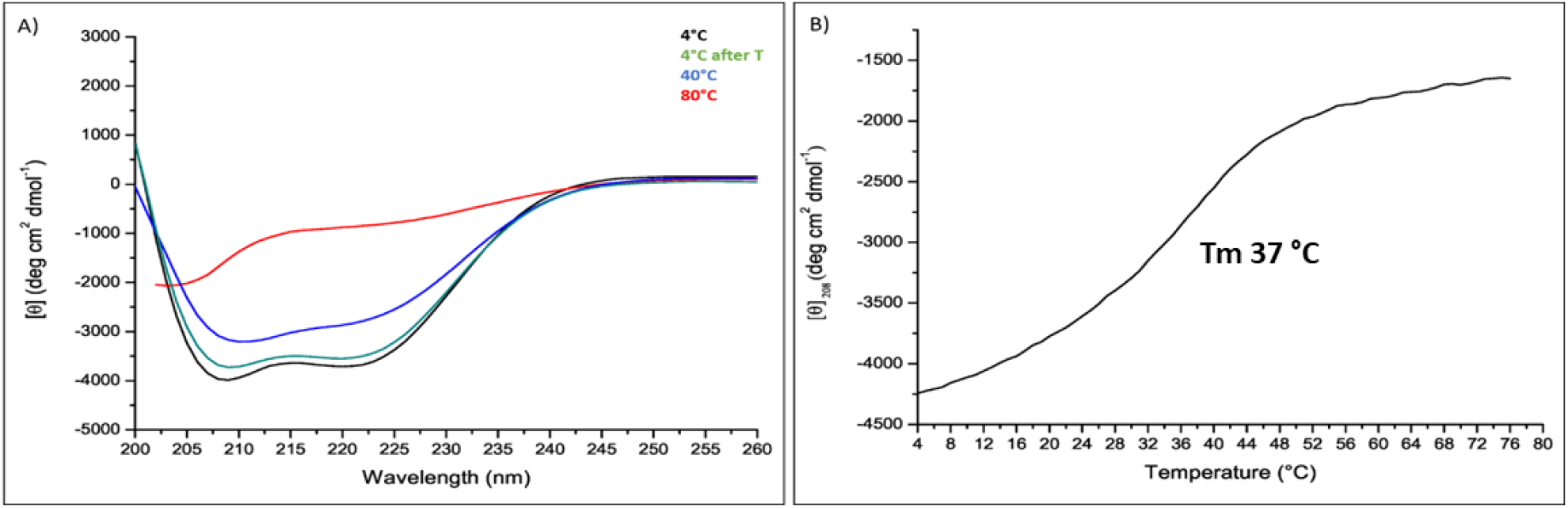
CD studies. (A) CD spectra of Spikeplug measured at 0.2 mg mL^-1^ in 20 mM NaP pH 7.4. (B) Thermal denaturation curve.

**Figure 4.**
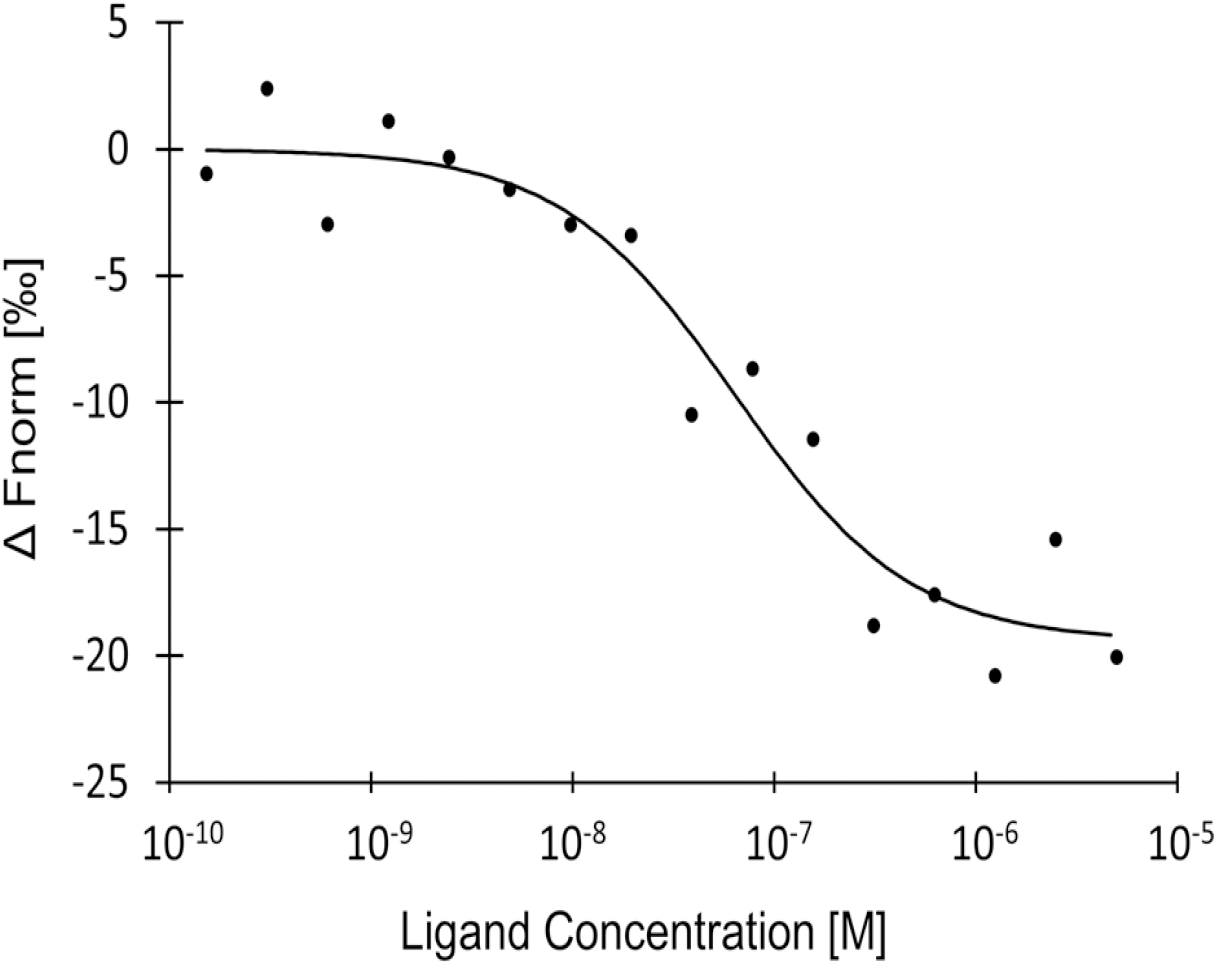
MicroScale Thermophoresis binding analysis for the interaction between Spike RBD and Spikeplug. The concentration of NT647-Spike RBD is kept constant at 25 nM, while the ligand concentration varies from 5 µM to 0.15 nM. The serial titrations result in measurable changes in the fluorescence signal within a temperature gradient that can be used to calculate the dissociation constant (Kd = 40 ± 5 nM). The curve is shown as ΔFnorm (change of Fnorm with respect to the zero ligand concentration) against Spikeplug concentration on a log scale.

## Discussion

Blocking the very first step of SARS-CoV-2 entry into host cells by hampering the interaction of the S protein with the cellular receptor, represents a highly promising therapeutic strategy against COVID-19. We focused on generating a Spike interactor that resembles the human ACE2 receptor as we argued that such type of molecule would act against all known coronaviruses using this receptor, such as SARS-CoV and HCoV-NL63, as well as other coronavirues possibily emerging in the future. Several structures of the complex between the S protein and the ACE2 receptor have been recently published. Based on this structural information, we designed a mini-protein which embeds all important Spike-interacting residues of the ACE2 receptor. This miniaturised mimic of ACE2, was properly engineered to be stable in solution and to enhance its affinity with the S protein. Indeed, an important issue with peptide- or protein-based drugs is its conformational stability. This mini-protein, Spikeplug, presents a highly stable α helical conformation in solution and a nanomolar binding affinity to the RBD domain of glycosylated S protein from SARS-CoV-2. The dissociation constant K_D_ determined here using MST, 40 nM, is in the same range of the affinity measured between the S protein and the ACE2 receptor of 14.7 nM [5]. We are currently designing additional variants to further improve binding affinity to the S protein.

Our basic idea is that mimics of the host receptor have a great advantage over other drugs for several reasons. First, if a novel coronavirus will emerge, able to infect humans through the ACE2 receptor, then its S proteins will be recognised and plugged by our receptor mimics (Figure 5), blocking viral entry. Importantly, Spikeplug is large enough to cover most important interactions with the S protein, but small enough to bind simultaneously to the three chains of the S protein, thus increasing its potential to hamper interactions with the ACE2 receptor (Figure 5). Another point we considered in our research was drug resistance, as pathogens can easily mutate to escape therapeutic treatments. However, if the virus mutates to escape binding to Spikeplug, then it will also likely drop its affinity for the natural ACE2 receptor, resulting in suicide action. The validity of such an approach has been demonstrated by the development of the antiretroviral drug Enfuvirtide, which blocks the action of the HIV-1 gp41 fusion protein, thus preventing viral entry. As for all antiviral drugs, also Enfuvirtide can select for resistance mutations. However, it has been shown that resistance to the drug comes at a serious fitness cost for the virus since it involves structural alterations in an essential component of the virus-host cell fusion complex [13]. Hence, targeting the receptor binding domain of SARS-CoV-2 might also result in a high genetic barrier towards resistance. Last, we expect Spikeplug to be well tolerated because of its almost complete identity with the correspondent domains of the endogenous human protein.

**Figure 5.**
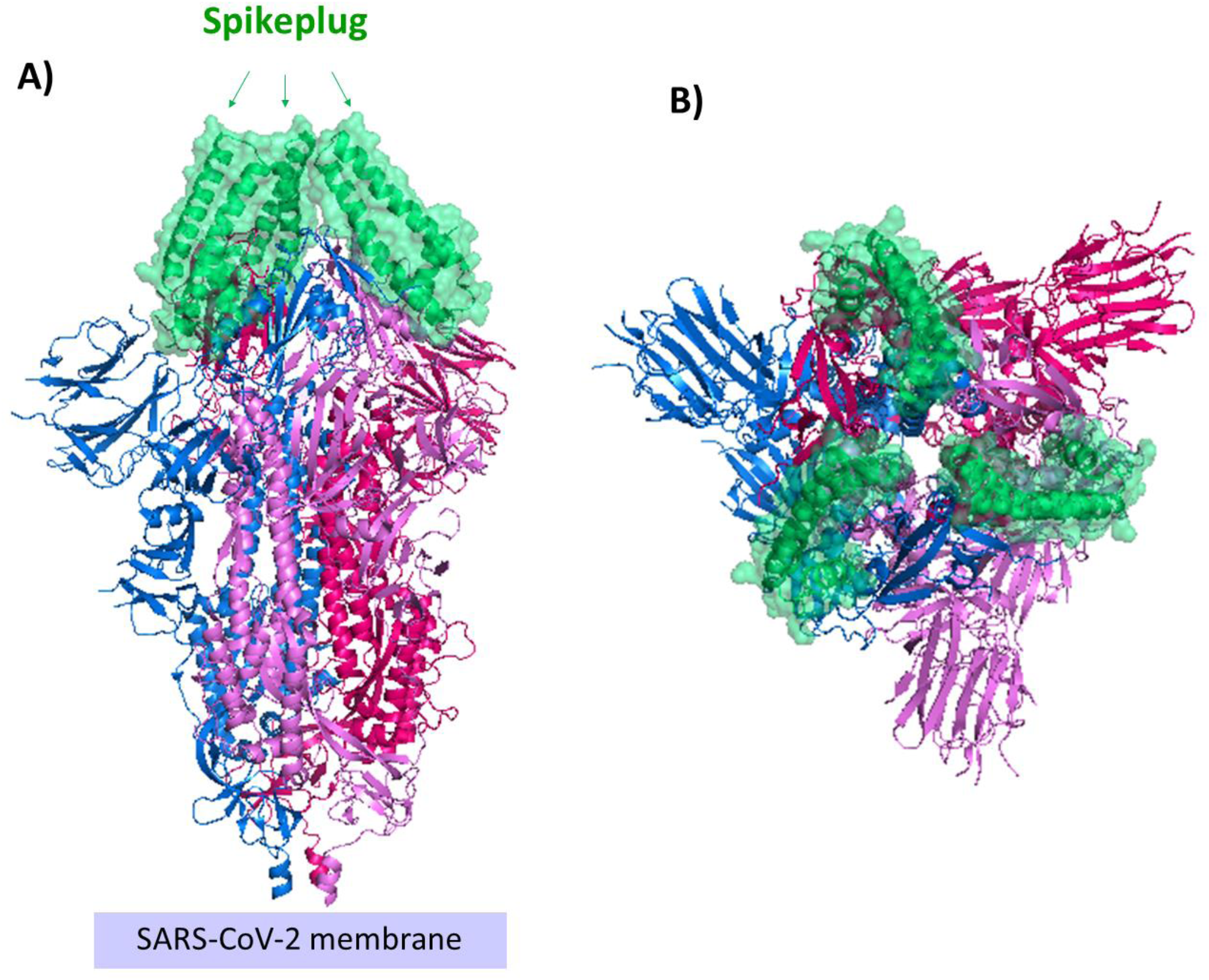
Cartoon representation of a plugged Spike trimer. The structure of the S protein (pdb code 6vxx) [9] is reported in magenta, prune and blue. The position of Spikeplug molecules (green surface representation) on the S protein was determined upon superposition of RBD domain of our minimised Spikeplug-RBD complex on the RBD domain of the entire S protein (pdb code 6vxx). A and B panels report side and top views, respectively.

In conclusion, we believe that Spikeplug is a first-in class promising lead candidate for the development of molecular targeted therapies against SARS-CoV-2 and other coronaviruses.

## Experimental methods

### Molecular design

Molecular modelling was initially performed using the cryo EM structure of the SARS-CoV-2 S protein (PDB code 6vsb) [5] and the crystal structure of the complex between ACE2 and the RBD domain of SARS-CoV S protein (PDB code 2ajf) [11]. To complete missing regions in the cryo EM structure, we computed the homology model of SARS-CoV-2 S protein using MODELLER [14], and the structure 2ajf as a template. While this work was in progress, the crystal structure of the complex between ACE2 and the RBD domain of the S protein from SARS-CoV-2 (PDB code 6m0j) was released [10]. In this structure, most of the interactions between the S protein and ACE2 are conserved. Given the high sequence identity between RBD of SARS-CoV and SARS-CoV-2 (84%), we relied on both structures to take also possible crystal packing bias into account. Mutations in the basic scaffold were generated using the software Coot [15]. Models were energy minimised using the GROMACS package [12]. Figures were generated with Pymol [16].

### Protein expression and purification

The gene encoding Spikeplug was synthesised and codon-optimised for *E*. *coli* expression by GeneArt® Gene Synthesis company (Invitrogen) and subsequently sub-cloned into pETM-13 vector (EMBL, Germany) between the NcoI and XhoI restriction sites. This plasmid endows the placement of a histidine-tag at the C-terminal of the mini-protein. The expression level was optimised after carrying out small scale expression screening using different *E*. *coli* available strains. From the screenings, the recombinant protein was successfully over-expressed in *E*.*coli* BL21(DE3) cells, resulting in a yield of 70 mg of pure protein from one litre of bacterial culture. Briefly, an overnight starting culture of 10 mL was prepared for growth in 1L of Luria Bertani (LB) medium containing 50µg L^−1^ kanamycin, which was then induced with 0.8 mM of IPTG at 16°C for approximately 16 h. The protein was purified by sonicating bacterial cells resuspended in binding buffer (300mM NaCl, 50mM Tris-HCl, 10 mM imidazole, 2.5% (v/v) glycerol, pH 7.8) containing a protease-inhibitor cocktail (Roche Diagnostics). The lysate was cleared by centrifugation at 18000 rpm at 4°C and the supernatant was loaded onto 5 mL Ni–NTA resin (Qiagen, Milan, Italy) equilibrated with binding buffer. After washing with ten volumes of binding buffer, the protein was eluted by adding 300 mM imidazole to the binding buffer. The fractions containing the eluted protein were pooled, concentrated and then loaded onto a Superdex 75 HR 10/30 gel-filtration column (GE Healthcare) equilibrated with 150mM NaCl, 50mM Tris-HCl, 2.5% (v/v) glycerol pH 7.8 for a further purification step. The sample eluted in a single peak and was homogeneous, as judged by 18% SDS–PAGE analysis. The protein was concentrated using a centrifugal filter (Merck Millipore) and the concentration was determined by UV absorbance using the corresponding ε values (M^-1^ cm^-1^).

### CD spectroscopy

CD measurements were carried out using a JASCO J-815 CD spectropolarimeter equipped with a Peltier temperature controller (Model PTC-423S) at 4°C in a 0.1 cm optical path length cell in the 200–260 nm wavelength range. Protein concentration was 0.2 mg/mL in 20 mM sodium phosphate buffer (pH 7.4). The data was recorded with a scanning speed of 20 nm/min and a band width of 1 nm. All spectra were averaged from three scans and baseline-corrected using a blank consisting of the protein buffer. The molar ellipticity per mean residue, [Ѳ] in deg·cm^2^·dmol−^1^, was calculated from the following equation: [Ѳ] = [Ѳ]obs × mrw × (10 × l × C)−^1^ where [Ѳ]obs is the ellipticity measured in degrees, mrw is the mean residue molecular mass (117.9 Da), l is the optical path length of the cell in cm, and C is the protein concentration in g/L. During the melting experiments, CD spectra were collected with increasing temperature from 4°C to 80 °C with an average rate of 1 °C/min. Thermal denaturation was investigated by recording the CD signal at 208 nm.

### Microscale thermophoresis (MST) binding studies

The thermophoretic measurements were performed using Monolith NT.115 device with red detection channel (NanoTemper Technologies, Munich, Germany). Recombinant Spikeplug was produced in our laboratory, while SARS-CoV-2 Spike RBD was purchased from Sino Biological Inc. For MST recording, the SARS-CoV-2 Spike RBD was labelled with the fluorescent dye NT647 using the protocol suggested from the NanoTemper. Thermophoretic experiments were conducted using Monolith NT.115 (NanoTemper Technologies, Munich, Germany). Briefly, 90 µl of a 10 µM solution of SARS-CoV-2 Spike protein (RBD) in labelling buffer (130 mM NaHCO_3_, 50 mM NaCl, pH 8.2) was mixed with 10 µl of 300 µM NT647-N-hydroxysuccinimide fluorophore (NanoTemper Technologies) and incubated in the dark for 30 min at room temperature. Spike RBD concentration after labelling (NT647-Spike RBD) was measured using a UV-Vis spectrophotometer and the labelling efficiency was determined to be about 25%.

The MST experiment was performed in a buffer containing 200 mM NaCl, 50mM Tris-HCl, 0.05% (v/v) Tween-20, 2.5% (v/v) glycerol, pH 7.8. 10 µL of NT647-Spike RBD 50 nM was mixed with 10 µL of 16 serial dilution of Spikeplug. The final concentration of NT647-Spike RBD was 25 nM in all samples, whereas Spikeplug concentration ranged from 0.15 nM to 5 μM. Samples were then loaded into sixteen premium-coated capillaries (NanoTemper Technologies) and fluorescence was recorded for 20 s using 100% laser power and 40% MST power. The temperature of the instrument was set to 25°C for all measurements. After recording the MST time traces, data were analysed. K_D_ value was calculated from ligand concentration-dependent changes in the normalised fluorescence (F_norm_) of NT647-Spike RBD after 14 s of thermophoresis. The assay was performed in duplicate and the values reported were generated through the usage of MO Affinity Analysis software (NanoTemper Technologies).

## Abbreviations

COVID-19: Coronavirus disease 19
PDB: Protein Databank
RBD: Receptor Binding Domain
RBM: Receptor Binding Motif
ACE2: angiotensin-converting enzyme 2
MST: Microscale thermophoresis.

## Acknowledgements

We thank Prof. Giovanni Maga, Institute of Molecular Genetics IGM-CNR, for careful reading of this manuscript.

## Notes

### Competing Interest Statement

The authors have declared no competing interest.

